# Pan-cancer analysis of patient tumor single-cell transcriptomes identifies promising selective and safe CAR targets in head and neck cancer

**DOI:** 10.1101/2021.09.29.462485

**Authors:** Sanna Madan, Sanju Sinha, Silvio J. Gutkind, Ezra E. W. Cohen, Alejandro A. Schäffer, Eytan Ruppin

**Author notes:** To whom correspondence should be addressed at or.

## Abstract

**BACKGROUND:** Chimeric antigen receptor (CAR) T cell therapies have yielded transformative clinical success for patients with blood tumors, but their full potential remains to be unleashed against solid tumors. One challenge is finding selective targets: cell surface proteins that are expressed widely by cancer cells and minimally by healthy cells in the tumor microenvironment and other normal tissues.

**METHODS:** Analyzing pan-cancer patient tumor single cell transcriptomics data, we first define and quantify selectivity and safety scores of existing CAR targets for indications in which they are in clinical trials or approved. Selectivity scores are computed by the ability of a given surfaceome gene to classify tumor from nontumor cells in the tumor microenvironment. Safety scores are computed by mining healthy tissue transcriptomics and proteomics atlas data. Second, we identify new candidate cell surface CAR targets that have better selectivity and safety scores than the leading targets among those currently being tested, in an indication-specific manner.

**RESULTS:** Remarkably, in almost all cancer types, we cannot find such better targets, testifying to the overall near optimality of the current target space. However, in HPV-negative head and neck squamous cell carcinoma (HNSC), for which there is currently a dearth of existing CAR targets, we find five new targets that have both superior selectivity and safety scores. Among the HNSC new targets, we find a few that additionally are strongly essentiality in HNSC cell lines.

---

**CONCLUSIONS:** The newly identified cell surface targets lay a basis for future investigations of better HNSC CAR treatments.

## Background

Chimeric antigen receptor T cell (CAR-T) therapy has revolutionized cancer treatment. After a patient’s T cells are extracted, they are modified to recognize a tumor-specific protein via the addition of a synthetic CAR on the cell membrane, and then infused back into the patient. CAR-T treatment has resulted in durable remissions in patients, particularly in those with B-cell malignancies. Recently, CARs directed against the B-cell marker CD19 have been declared curative, with observed remission rates as high as 90% [1]. The physiological phenomenon that makes B cell tumors more amenable to CAR-T therapy than other tumors is that one can kill all the B cells – both cancerous and not cancerous – and the patient is likely to survive, especially if treated with intravenous immunoglobulin (IVIG) to replace the missing antibodies.

As of September 2021, the Trialtrove repository of clinical trials [2] had recorded more than 1,500 clinical trials involving cells engineered to target various membrane surface proteins or other antigens in specific indications that have either taken place, are currently in progress, or are proposed/planned. Most of these trials involve modified CAR T cells and some involve NK cells; because our analysis applies to both T and NK cells, we use “CAR” without a suffix in most places. The CAR targets that are currently FDA-approved or are being tested include a variety of proteins, *e.g.*, CD19 and CD22 in B-cell malignancies, and HER2, IL-13 receptor α2, EGFRvIII, carbonic anhydrase IX, and MUC1 in various solid tumors [3].

While CAR-T treatments have been remarkably successful against B-cell tumors, several challenges remain for realizing their potential in the realm of solid tumors. Two key challenges are: (1) heterogeneity of cell surface proteins expressed by a tumor’s constituent malignant cells, and (2) expression of the CAR target protein on the cell surfaces of normal tissues elsewhere in the body, posing a toxicity risk. These challenges may be addressed by studying single-cell RNA sequencing (scRNA-seq), which has generated many patient tumor transcriptomes in recent years. We hypothesized that we could harness these single-cell measurements to profile the expression of genes encoding targets of CAR therapies and more broadly, cell surface proteins, at a cellular resolution – thus enabling us to identify suitable surface proteins that had not previously been considered seriously as targets for CAR therapy. Indeed, recent studies have harnessed scRNA-seq to investigate toxicities of CAR-T therapies [4,5] and highlighted the importance of optimizing surface target selection by utilizing scRNA-seq [6–8]. Furthermore, a notable recent analysis leveraged public patient tumor and reference atlas scRNA-seq data to first construct an integrated “meta-atlas” of tumor and normal cells and then identify tumor-specific combinations of target antigens via logic gates [9]. Hopefully, identifying optimal cell surface targets in a systematic, data-driven manner may translate in the future to improved patient outcomes, leading to more durable remissions and reduced toxicities.

Zeroing in on the challenges listed above, we asked whether mining scRNA-seq datasets of solid tumors can enable us to: (1) First, chart the landscape of existing cell-surface targets against which CARs are currently aimed at in clinical trials, and estimate their selectivity and safety with quantitative scores. (2) Second, identify alternative target proteins that are differentially highly expressed by tumor cells and lowly expressed by non-tumor cells within the tumor microenvironment (TME) and additionally in normal tissues across the body. Our goal is to identify new targets for CAR therapies that have better safety and selectivity scores than the targets currently tested, in an indication-specific manner.

We use the term “indication” to refer to an ordered pair of (cancer type, CAR target) that has been proposed in at least one clinical trial; we use the opposite term “non-indication” to denote a (cancer type, CAR target) for which we could not find any proposed clinical trial (**Methods**). Our analysis proceeds in the following steps: (1) First, we mine clinical trial databases and literature, we identify the current CAR-T targets in clinics mapped to the solid tumors in which they are indicated / targeted against. For each (cancer type, CAR target) pair, we compute a **tumor selectivity score**, *i.e.*, the selectivity of the expression of a given candidate target gene in tumor cells versus all other cells in the TME. This is quantified by the ability of a classifier to separate tumor cells from non-tumor cells based on the single cell expression data of this gene in the TME of the respective cancer indication. (2) Second, we compute a **safety score** for each candidate target, which is computed by quantifying the level of protein and scRNA-seq expression across normal human tissues. We additionally investigate the *in vitro* essentiality of genes encoding CAR targets that are either approved or being tested in clinics as well as the candidate new targets yielded by our analysis in CRISPR knockout screens. The overview of our approach showing all three resource inputs is shown in **Figure 1**.

**Figure 1:**
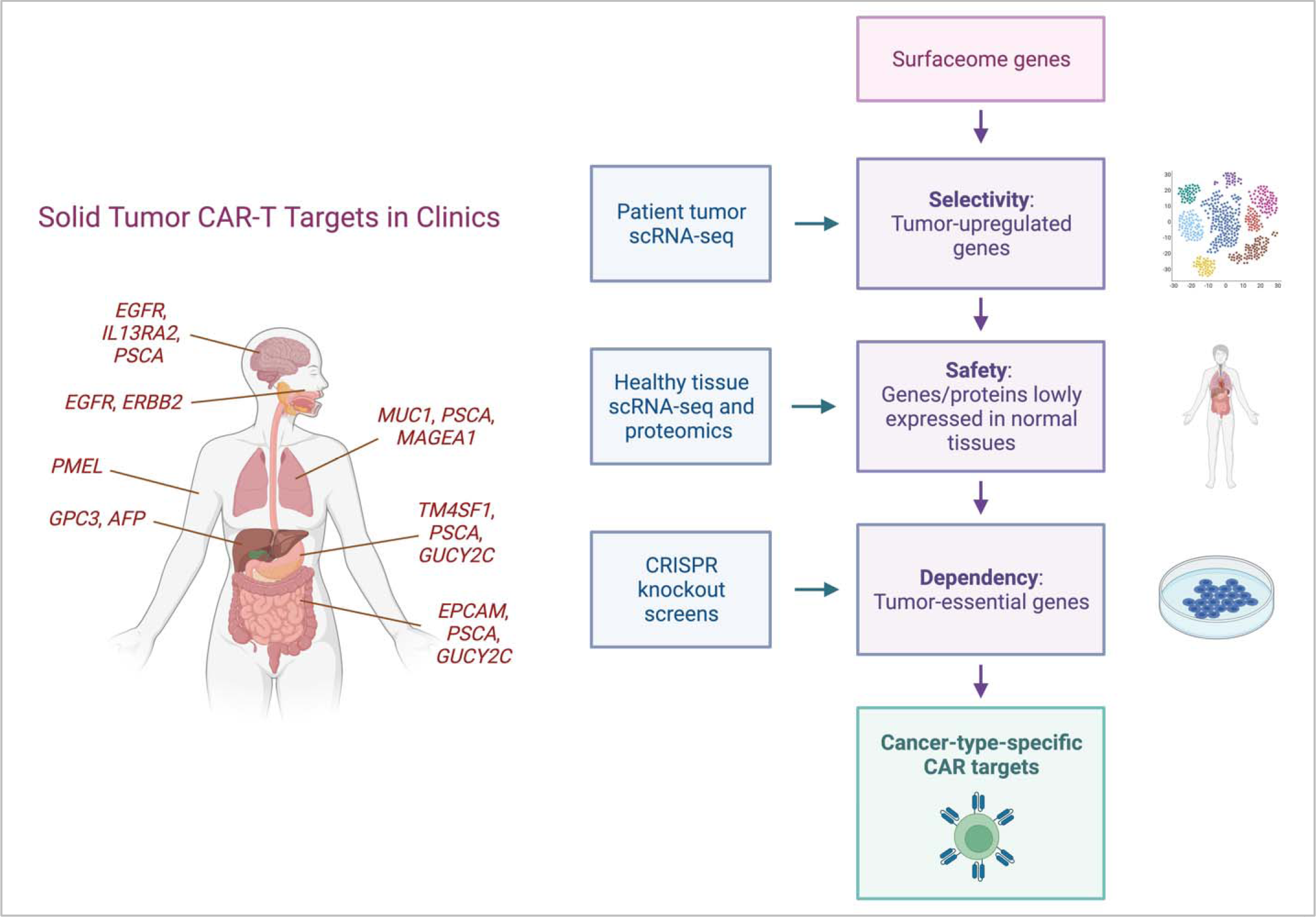
The left panel shows some solid cancer locations and CAR-T targets being tested for them. The right panel shows that our study uses single-cell RNA data from patients, healthy cell atlases, and CRISPR knockout screens as inputs to help us select CAR-T targets on the right.

The first part of our analysis evaluates the indications that are currently studied in clinical trials (or already approved) for their tumor selectivity and safety metrics. We then compare untested, candidate CAR target surface protein versus these current state-of-the-art benchmarks. Our goal is to identify new target genes that match or exceed the leading extant targets’ selectivity and safety scores in each respective cancer type, thus forming promising new target tumor antigens for further possible consideration when thinking about engineering new CARs. In difference from previous related studies [7,9], the unique aim of our study is to find novel targets for CARs that are strictly better than the best CAR target antigens in clinics based on the selectivity and safety metrics. This analysis, quite surprisingly, identifies only a few but highly promising candidates for clinical translation.

## Methods

### Clinical trial curation

To obtain a list of CAR clinical trials and targets in solid tumors, we first obtained a list of single clinical CAR targets from MacKay *et al.*, who carried out their ClinicalTrials.gov query on March 3, 2019 [10]. To obtain a complete up-to-date list of clinical trials since then, an additional search was carried out on ClinicalTrials.gov on August 20, 2021 for CAR-T clinical trials posted on March 3, 2019 and after. The following query was used: “*CAR-T OR CAR OR chimeric antigen receptor OR chimeric immunoreceptors OR artificial T-cell receptors | Recruiting, Not yet recruiting, Active, not recruiting, Completed, Enrolling by invitation, Suspended, Terminated Studies | Interventional Studies | (cancer OR carcinoma OR solid tumor OR solid tumors OR melanoma OR sarcoma) NOT (hematological OR lymphoid OR liquid cancers OR lymphoma OR leukemia OR CD19 OR B cell malignancy OR myeloma OR Hematologic OR blood cancer) | First posted from 03/03/2019 to 08/20/2021*”. CAR trials and targets were further identified through manual curation, and clinical trial statuses were cross-checked on TrialTrove (https://citeline.informa.com/trials/results) to ensure consistency [2]. Access to TrialTrove requires a license.

CAR protein targets were then selected for further analyses if they had a corresponding single gene that could be analyzed for gene expression. The following molecules were left out of the analysis: Claudin 18.2, CD44v6, EGFRvIII, k-IgG, LeY, CEA, and GD2 because i) they are not proteins or ii) they are protein isoforms whose presence and abundance cannot easily be estimated from single cell gene expression data. For each target, its indications were identified as the specific tumor types which were mentioned in its ClinicalTrials.gov records, excluding ‘solid tumors’ and ‘metastases’. The following mapping of clinical trial cancer types to our datasets was used: brain cancer or glioma → glioblastoma; pancreatic cancer → pancreatic adenocarcinoma; stomach cancer or gastric cancer → stomach adenocarcinoma; liver or hepatocellular carcinoma → liver hepatocellular carcinoma; head and neck cancer → head and neck squamous cell carcinoma; melanoma → skin cutaneous melanoma; colon cancer or colorectal cancer → colorectal cancer; ovarian cancer → ovarian serous cystadenocarcinoma; lung cancer or non-small cell lung cancer (but not small cell lung cancer) → non-small cell lung cancer; breast cancer → breast invasive carcinoma. Liquid tumor CAR targets were taken from MacKay *et al*. [10]

### Single-cell transcriptomics data collection

We downloaded single cell transcriptomics datasets from the Tumor Immune Single-cell Hub (TISCH) 1.0 database [11]. This resource contains 76 uniformly processed scRNA-seq datasets spanning 28 cancer types and containing data from over 2 million cells. We selected datasets from human tissues which contained *both* malignant and non-malignant cells. This yielded a total of 26 unique solid tumor datasets: two of breast cancer (BRCA), one of colorectal cancer (CRC), eleven of glioblastoma (GBM), one of head and neck squamous cell carcinoma (HNSC), one of liver hepatocellular carcinoma (LIHC), four of non-small cell lung cancer (NSCLC), one of ovarian cancer (OV), two of pancreatic adenocarcinoma (PAAD), two of skin cutaneous melanoma (SKCM), and one of stomach adenocarcinoma (STAD). Additionally, an acute myeloid leukemia (AML) dataset was analyzed. These data were all uniformly processed by TISCH as log_2_(TPM/10+1) values (TPM stands for transcripts per million).

### Single-cell data analysis

In-house programs written in Python 3 were used to carry out analyses on the scRNA-seq datasets. A cell was labeled ‘tumor’ if it was labeled ‘malignant’ by TISCH, and ‘non-tumor’ otherwise. We then partitioned the cells in each dataset by patient (if such annotation was available); the partition by patient is so that each patient counts equally in the downstream analysis regardless of how many cells were sampled from that patient. As quality control, only patient samples which contained at least 25 tumor cells and 25 non-tumor cells were selected for analysis. After this filtering step, only the cancer types where we had samples from at least three patients remaining were subsequently considered, excluding breast cancer entirely from our analysis, for which we only had two patients.

A list of 3,559 cell surface protein encoding genes was obtained from Hu *et al* [7]. Genes whose symbols differed in that reference in TISCH and in DepMap (see later subsection) were resolved by using the Human Gene Nomenclature Committee table including gene aliases (downloaded from https://www.genenames.org) We generated patient-specific-AUCs of tumor-vs-non-tumor cell discrimination for each gene whose respective protein was listed in a CAR clinical trial and for all cell surface protein encoding genes at large. For each existing (CAR target, indication) we compute a mean tumor selectivity score over all patient-specific selectivity scores. We took the top-ranking target in each cancer type by the mean tumor selectivity scores as the threshold to which we compared all other cell surface protein encoding genes. In doing these comparisons, we used the Wilcoxon signed rank one-sided test comparing matched patient-level AUCs, with Benjamini-Hochberg FDR p-value < 0.1 denoting statistical significance, repeating this procedure for each cancer type separately.

### Reference atlas analysis: computing safety scores

For computing the safety score of a cell surface protein in the Human Protein Atlas (HPA), an approach similar to that outlined in MacKay *et al.* [10] was taken. Individual tissue types were grouped into tissue groups, with the following mapping:

| **Tissue Group** | **Tissue types** |
| --- | --- |
| Adipose and soft | adipose tissue, soft tissue |
| Endocrine | adrenal gland, parathyroid gland, thymus,<br>thyroid gland |
| Nervous system | caudate, cerebellum, cerebral cortex, choroid |
|  | plexus, hippocampus, hypothalamus, pituitary gland, substantia nigra, dorsal raphe |
| Circulatory | heart muscle |
| Immune tissue | appendix, lymph node, spleen, tonsil |
| Immune cells | bone marrow |
| Female tissue | breast, cervix, endometrium, fallopian tube, lactating breast, ovary, placenta, vagina |
| Lungs | bronchus, lung, nasopharynx |
| Colon | colon |
| Gastrointestinal | duodenum, oral mucosa, rectum |
| Male tissue | epididymis, prostate, seminal vesicle, testis |
| Esophagus | esophagus |
| Eye | eye, retina |
| Gallbladder | gallbladder |
| Skin | hair, skin, sole of foot |
| Urinary | kidney, urinary bladder |
| Liver | liver |
| Pancreas | pancreas |
| Salivary gland | salivary gland |
| Muscle | skeletal muscle, smooth muscle |
| Small intestine | small intestine |
| Stomach | stomach |

HPA protein measurements for a given tissue type are either ‘Not detected’, ‘Low’, ‘Medium’, or ‘High’. For a tissue group, if the measurements of a protein over all tissue types only ranged from ‘Not detected’ to ‘Low’, then the tissue group was assigned ‘ND-L’. If the measurements over all tissue types only ranged from ‘Medium’ to ‘High’, then the group was assigned ‘M-H’. If values from either ‘Not detected’ or ‘Low’ AND ‘Medium’ or ‘High’ were present in the measurements over all tissue types, then the group was assigned ‘Variable.’ Thus, in determining the overall HPA safety score of a protein, the scoring weight is in the following order: ND-L > Variable > M-H. For each tissue group, a score of 2 was added for each ‘ND-L’, 1 was added for each ‘Variable’, and 0 was added for each ‘M-H.’ The mean of the resulting sum was taken to obtain a representative measure over all tissue groups that were measured for the protein. As such, the scoring values ranged from 0 to 2.

For the Tabula Sapiens safety score computation, the tissues measured were taken as is, with no further grouping, as the following: Bladder, Blood, Bone Marrow, Eye, Fat, Heart, Kidney, Large intestine, Liver, Lung, Lymph node, Mammary Gland, Muscle, Pancreas, Prostate, Salivary Gland, Skin, Small intestine, Spleen, Thymus, Tongue, Trachea, Uterus, and Vasculature. To compute the safety score of a surfaceome gene, first, within each tissue, the percentage of cells expressing that gene was computed. Next, for a tissue, if the percentage was < 1%, 10 was added to the final score; if it fell between 1-10%, 9 was added to the final score; if it fell between 11-20%, 8 was added to the final score; if it fell between 21-30%, 7 was added to the final score; if it fell between 31-40%, 6 was added to the final score; if it fell between 41-50%, 5 was added to the final score; if it fell between 51-60%, 4 was added to the final score; if it fell between 61%-70%, 3 was added to the final score; if it fell between 71-80%, 2 was added to the final score; if it fell between 81-90%, 1 was added to the final score; and finally, if it fell between 91-100%, 0 was added to the final score. The mean of this sum was taken over the total number of tissue groups for which the gene had been measured. As such, the scoring values ranged from 0 to 10.

### DepMap essentiality scores

We used the DepMap Public 22Q2, specifically ‘CRISPR_gene_dependency.csv’ was downloaded from https://depmap.org/portal/. This file contains the probability of dependence scores for each (cell line,gene) pair. The DepMap defines a probability of ≥ 0.5 as essential, and as such, our analysis defined a cell line as dependent on a gene if its *p*(dependent) ≥ 0.5.

Cell lines were partitioned by cancer type for each clinical or novel CAR target gene. For CRC, we analyzed cell lines labeled ‘Colon/Colorectal Cancer’ as the primary disease; for GBM, we analyzed cell lines labeled ‘Brain Cancer’ as primary disease and ‘Glioblastoma’ as the subtype; for NSCLC, we analyzed cell lines labeled ‘Lung Cancer’ as primary disease and ‘Non-Small Cell Lung Cancer (NSCLC)’ as the subtype; for PAAD we analyzed cell lines labeled ‘Pancreatic cancer’ as the primary disease and ‘Adenocarcinoma’ as the subtype; for STAD we analyzed cell lines labeled ‘Gastric Cancer’ as the primary disease and ‘Adenocarcinoma’ as the subtype; for OV we analyzed cell lines labeled ‘Ovarian Cancer’ as the primary disease with no constraint on subtype; for SKCM we analyzed cell lines labeled ‘Skin Cancer’ with ‘Melanoma’ as the subtype; for HNSC we analyzed cell lines labeled ‘Head and Neck Cancer’ as the primary disease and ‘Squamous Cell Carcinoma’ as the subtype; for B-cell Non-Hodgkin’s Lymphoma we analyzed cell lines labeled ‘Lymphoma’ as the primary disease and ‘B-cell, Non-Hodgkin’s’ as the subtype; for B-cell Hodgkin’s Lymphoma we analyzed cell lines as primary ‘Lymphoma’ and subtype ‘B-cell, Hodgkin’s’; for T-cell, Non-Hodgkin’s Lymphoma, we analyzed cells labeled ‘Lymphoma’ as primary disease and ‘T-cell, Non-Hodgkin’s’ as subtype; for T-cell Lymphoma we analyzed cell lines with ‘Lymphoma’ primary and ‘T-cell’ subtype; for B-cell Lymphoma, we analyzed cell lines with ‘Lymphoma’ primary and ‘B-cell’ subtype; for Acute Myelogenous Leukemia (AML) we analyzed cell lines with ‘Leukemia’ primary and ‘Acute Myelogenous Leukemia’ in the subtype; for Chronic Myelogenous Leukemia we analyzed cell lines with ‘Leukemia’ as the primary and ‘Chronic Myelogenous Leukemia’ as the subtype; for B-cell Chronic Lymphoblastic Leukemia (B-CLL), we analyzed cell lines with ‘Leukemia’ as the primary and ‘Chronic Lymphoblastic Leukemia’ as the subtype; for B-cell Acute Myeloid Leukemia (AML) we analyzed cell lines with ‘Leukemia’ as the primary and ‘Acute Lymphoblastic Leukemia (ALL), B-cell’ as the subtype; for T-cell AML, we analyzed cell lines with ‘Leukemia’ as the primary and ‘Acute Lymphoblastic Leukemia (ALL), T-cell’ as the subtype; for unspecified B-cell Leukemia we analyzed cell lines with ‘Leukemia’ as the primary and ‘B-cell, unspecified’ as the subtype; for Myeloma we analyzed cell lines with ‘Myeloma’ as the primary.

For each (cancer type, gene) indication, the mean *p*(essentiality) of the gene was simply taken over all *p*(essential) values in the cell lines of the cancer type indication as mapped above.

The DepMap (https://depmap.org/portal/documentation/) defines a ‘strongly selective’ gene “as having a dependency distribution that is at least 100 times more likely to have been sampled from a skewed t-distribution than a normal distribution.” *i.e.*, its skewed-LRT value exceeds 100. A ‘common essential’ (*i.e.*, ‘pan-essential’) gene “is defined as a gene which, in a large, pan-cancer screen, ranks in the top X most depleting genes in at least 90% of cell lines. X is chosen empirically using the minimum of the distribution of gene ranks in the 90th percentile least depleting lines.”

## Results

### Overview of the analysis

We analyzed nine cancer types that have sufficient data in the scRNA-seq Tumor Immune Single-cell Hub (TISCH) 1.0 collection: colorectal cancer, glioma, (HPV-negative) head and neck cancer, liver cancer, non-small cell lung cancer, ovarian cancer, pancreatic adenocarcinoma, melanoma, and stomach cancer [11] (**Methods**). To quantify the *tumor selectivity* of a given gene encoding either an extant or new candidate CAR target cell surface protein, we used the “area under the curve” (AUC) metric, which quantifies the extent by which its expression levels can be used to discriminate between tumor and non-tumor cells in the TME (**Methods**). We use ‘AUC’ and ‘*tumor selectivity score*’ interchangeably. The tumor selectivity score is computed for each patient individually, such that for a given cell surface protein target in a cancer type, we obtain a distribution of tumor selectivity scores across all patients, whose mean and standard deviation can be quantified.

In parallel, to quantify the *overall safety* of a target gene encoding a cell surface protein, a two-part *safety score* was computed, considering separately the Human Protein Atlas (HPA) proteomics data [12] and the scRNA-seq Tabula sapiens gene expression data [13]. Each of these two *sub-safety scores* is computed at the tissue level and averaged over the set of pertaining samples; for each resource, the safety measures are computed for each tissue separately and then the arithmetic mean of safety scores across all tissues is taken as the final safety score (**Methods**). The HPA safety scores range from 0-2, and the Tabula Sapiens safety scores range from 0-10, following each resource’s scoring.

### The quantified tumor selectivity and safety landscape of the human surfaceome

To obtain a birds-eye view of the tumor selectivity and safety metrics outlined in the previous section, we quantified their values across all surfaceome genes (curated from Hu *et al*. [7]). The cancer-type-specific distributions of tumor selectivity scores across all surfaceome genes are shown in **Figure 2A**. The distributions are highly similar across cancer types: all are symmetric and unimodal with AUC values tightly concentrated around 0.50.

**Figure 2:**
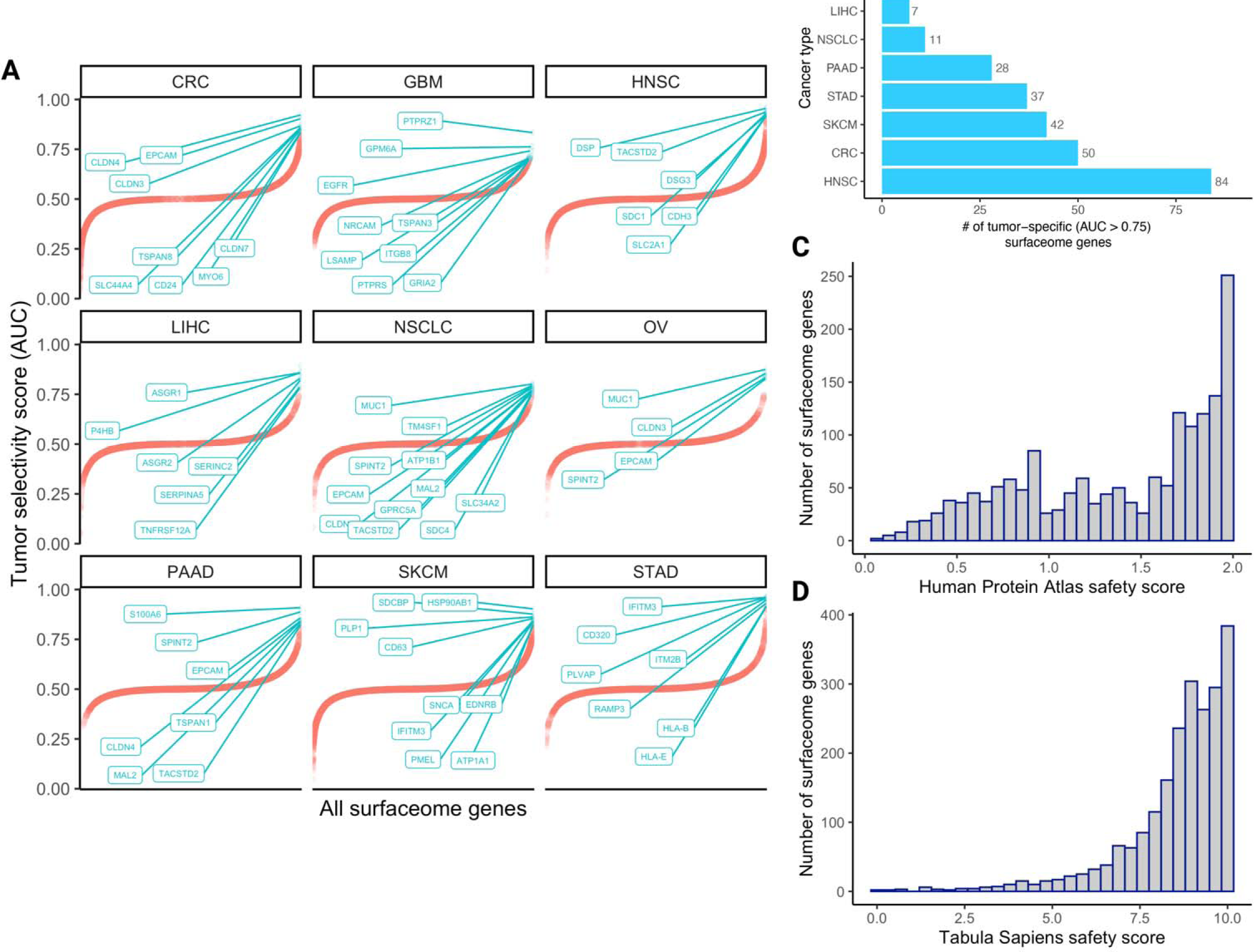
The landscape of tumor selectivity and safety scores of human surfaceome genes. **(A)** 1D scatterplots depicting the tumor selectivity scores (AUCs) of all surfaceome genes within each cancer type (red curves). Within each cancer-type-plot, the top N (where N is < 10) genes are labeled and represented by green points instead of red. Genes are sorted in ascending order on the x-axis from left to right. **(B)** The number of tumor-specific surfaceome genes (AUC > 0.75) in each cancer type, in ascending order from top to bottom. Histograms depicting the distributions of HPA-derived safety scores and Tabula sapiens-derived safety scores are shown in **(C)** and **(D)**, respectively.

We examined how many highly tumor-specific (AUC > 0.75) surfaceome genes were upregulated in each cancer type (**Figure 2B**). Interestingly, the number of such surfaceome genes varies considerably by cancer type, with the lowest values in GBM (n=2), OV (n=6), and LIHC (n=7), and highest values in SKCM (n=42), CRC (n=50), and HNSC (n=84). This suggests that the latter subset of solid tumor types may contain a larger space of surface targets that may be considered for CAR therapeutic development, compared to the former subset of solid tumor types. Finally, the distribution of safety scores derived from the HPA and Tabula sapiens atlases are shown in **Figure 2C** and **Figure 2D**, respectively. Both distributions are strongly skewed to the left; most surfaceome genes and proteins have considerably high safety scores (HPA safety mean=1.38, median=1.55, standard deviation=0.53; Tabula sapiens safety mean=8.56, median=9, standard deviation=1.54). Additionally, the quantified cancer-type-specific selectivity scores and safety scores derived from each resource (with tissue-specific resolution) for all surfaceome genes are provided in **Additional File 2: Tables S1-S3**.

### The selectivity and safety landscape of targets of approved and currently studied CARs

Mining TrialTrove, ClinicalTrials.gov, and existing literature [10], we assembled a table summarizing the genes encoding proteins that are currently being targeted by CARs in clinical trials in each cancer type, including those already approved **(Table 1 & Methods)**. We term those *clinical* CAR targets. For almost all the solid tumors we have considered, we find at least 10-20 unique genes encoding cell surface proteins that are currently targeted by CARs in various stages of active or planned clinical trials. The most frequently studied genes among those are *PSCA, MUC1, CD274, EGFR*, and *ERBB2*. Notably, however, head and neck cancer (HNSC) has a dearth of unique CAR targets, with only two targets currently in clinical trials, *EGFR* and *ERBB2*, suggesting that there may be a critical unmet need to identify additional cell surface targets for HNSC. Following this assessment, we charted the distribution of tumor selectivity scores for each cancer type’s pertaining clinical CAR targets (**Figure 3A, 3B**).

**Table 1:** Current solid tumor CAR targets in clinics. Genes encoding cell surface proteins that are currently targeted by CARs in clinical trials, for the nine solid tumor types we studied.

| Cancer type | Genes encoding clinical CAR targets |
| --- | --- |
| CRC | *MUC1, CD274, PROM1, ROR1, PSCA, MET, EPCAM, EGFR, FOLH1, ERBB2, MSLN, CD276, TM4SF1, CEACAM5, GUCY2C* |
| GBM | *MUC1, CD274, PROM1, ROR1, MPP2, EPHA2, PSCA, IL13RA2, EGFR, ERBB2, MSLN, CD276, BSG* |
| LIHC | *MUC1, CD274, PROM1, ROR1, AFP, TNFRSF10B, PTGS2, PSCA, MET, EPCAM, EGFR, GPC3, ERBB2, MSLN, CD276, BSG* |
| NSCLC | *MUC1, CD274, ROR1, CD80, CD86, MAGEA1, MAGEA4, PSCA, EGFR, FOLH1, GPC3, ERBB2, MSLN, CD276, TM4SF1* |
| PAAD | *MUC1, CD274, PROM1, CD70, ROR2, CLDN18, PSCA, EPCAM, EGFR, ERBB2, MSLN, TM4SF1, GUCY2C* |
| STAD | *MUC1, ROR2, PSCA, EPCAM, EGFR, ERBB2, MSLN, CD276, TM4SF1, GUCY2C* |
| OV | *PROM1, CD70, FOLR1, MUC16, MET, EGFR, ERBB2, MSLN, ALPP* |
| SKCM | *CD70, PMEL, KDR, MET, IL13RA2* |
| HNSC | *EGFR, ERBB2* |

**Figure 3:**
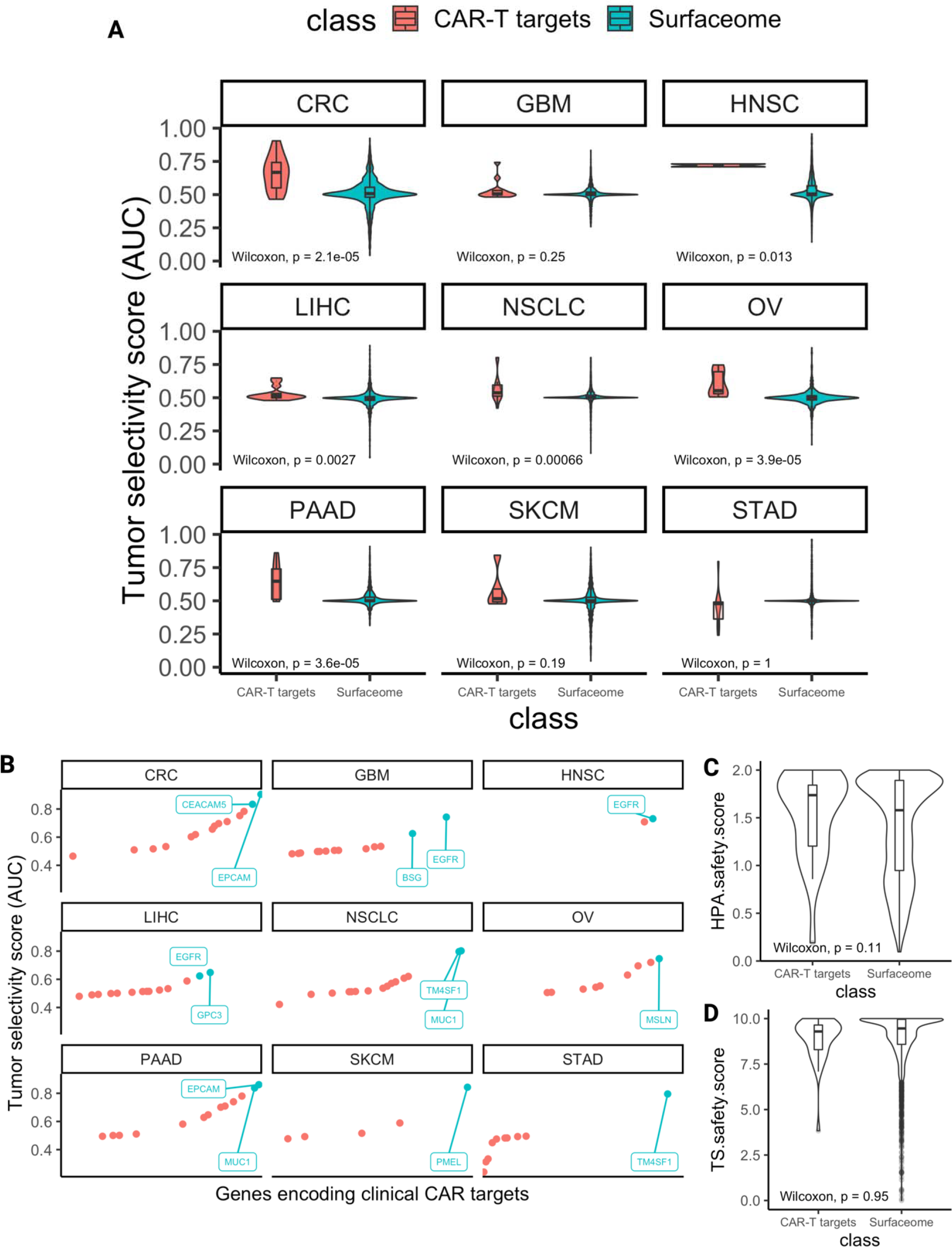
Surveying the selectivity and safety scores of known CAR targets. **(A)** Violin plots with nested boxplots depicting the tumor selectivity scores of genes encoding clinical CAR-T targets (shown in red) versus all surfaceome genes (shown in green) for each cancer type. Wilcoxon rank-sum test with p-values shown in each subplot, p < 0.05 significant. **(B)** 1D scatterplots depicting tumor selectivity scores (AUCs) of genes encoding clinical CAR targets within each cancer type. Genes whose selectivity scores are in the top 10% are represented by green points and are labeled, for each cancer type-plot. Genes are sorted in ascending order on the x-axis by AUC from left to right. **(C-D)** Distribution of safety scores of all clinical CAR target encoding genes versus all surfaceome genes. Tabula sapiens derived safety scores **(C)** and HPA derived safety scores **(D)**. Wilcoxon rank-sum comparison test p-values shown at the bottom of each plot.

Next, for each cancer type studied, we identified: 1) the leading clinical CAR target based on its tumor selectivity score, as derived from TISCH scRNA-seq data, 2) the leading clinical CAR target based on its safety score, as derived from the Human Protein Atlas (HPA), and 3) the leading clinical CAR target based on its safety score, as derived from the Tabula sapiens (TS) scRNA-seq reference atlas (**Table 2**). The scores of the leading targets by each criterion are further used as threshold parameters by which any gene encoding cell surface proteins can be compared to quantify its potential to serve as a CAR targets.

**Table 2:** The top selectivity and safety scores of current clinical CAR targets within each cancer type.

| Cancer type | Gene with highest tumor selectivity score (0-1) | Gene with highest HPA Safety score (0-2) | Gene with highest TS safety score (0-10) |
| --- | --- | --- | --- |
| STAD | *TM4SF1*<br>(0.80) | *PSCA*<br>(2.0) | *GUCY2C*<br>(9.83) |
| HNSC | *EGFR*<br>(0.73) | *ERBB2*<br>(1.55) | *ERBB2*<br>(7.92) |
| OV | *MSLN*<br>(0.75) | *MUC16*<br>(1.84) | *ALPP*<br>(10.0) |
| SKCM | *PMEL*<br>(0.84) | *PMEL*<br>(1.95) | *PMEL*<br>(9.79) |
| CRC | *EPCAM*<br>(0.90) | *PSCA*<br>(2.0) | *GUCY2C*<br>(9.83) |
| GBM | *EGFR*<br>(0.74) | *PSCA*<br>(2.0) | *IL13RA2*<br>(9.79) |
| PAAD | *EPCAM*<br>(0.86) | *PSCA*<br>(2.0) | *CLDN18*<br>(9.83) |
| LIHC | *GPC3*<br>(0.65) | *AFP*<br>(2.0) | *AFP*<br>(9.79) |
| NSCLC | *MUC1*<br>(0.80) | *PSCA*<br>(2.0) | *MAGEA1*<br>(10.0) |

We find that the tumor selectivity scores of genes encoding clinical CAR targets is higher than other surfaceome genes in a statistically significant manner in six cancer types (Wilcoxon rank-sum p-value < 0.05) (**Figure 3A**). Notably, this testifies to the utility of the single-cell tumor selectivity metric we have derived, as this measure is demonstrably higher among genes encoding surface proteins that have already been vetted to be targeted by CARs. We similarly chart the distributions of safety score values, also encouragingly revealing that the safety scores we derived from the HPA are higher among clinical CAR targets than all surfaceome proteins (**Figure 3C**, Wilcoxon rank-sum p=0.11). However, we note that the median Tabula sapiens-derived safety scores of all surfaceome genes and genes encoding clinical CAR targets are roughly equivalent (**Figure 3D**, Wilcoxon rank-sum p=0.95). We continue to use this sub-safety score in what follows to adopt a cautious approach and provide as comprehensive a picture as possible, but we note that it is not discriminatory as the tumor selectivity and HPA safety scores are. We additionally plotted the distributions of safety scores in all surfaceome genes and cancer-type-stratified clinical CAR targets (**Additional File 1: Figure S1**), as well as comparisons between genes encoding clinical CAR targets and the most tumor-selective surfaceome genes in each cancer type (**Additional File 1: Figure S2**).

### Identifying novel CAR targets with higher estimated selectivity and safety scores

In the previous subsection, for each cancer type, we identified the top-ranking *existing* CAR targets based on their tumor selectivity and safety scores. We now pivot to identify new candidate targets that have better selectivity and safety scores than those of the current therapies. We proceeded in three steps: (1) identifying highly selective CAR targets (with yet moderate toxicity levels); (2) identifying high safety scoring CAR targets (with yet moderate selectivity levels), and finally, (3) identifying the targets that have both very high selectivity and safety scores (**Figure 4A**). These three steps were conducted for every indication, comparing the scores of the new targets to the best scoring CAR targets in that cancer type.

**Figure 4:**
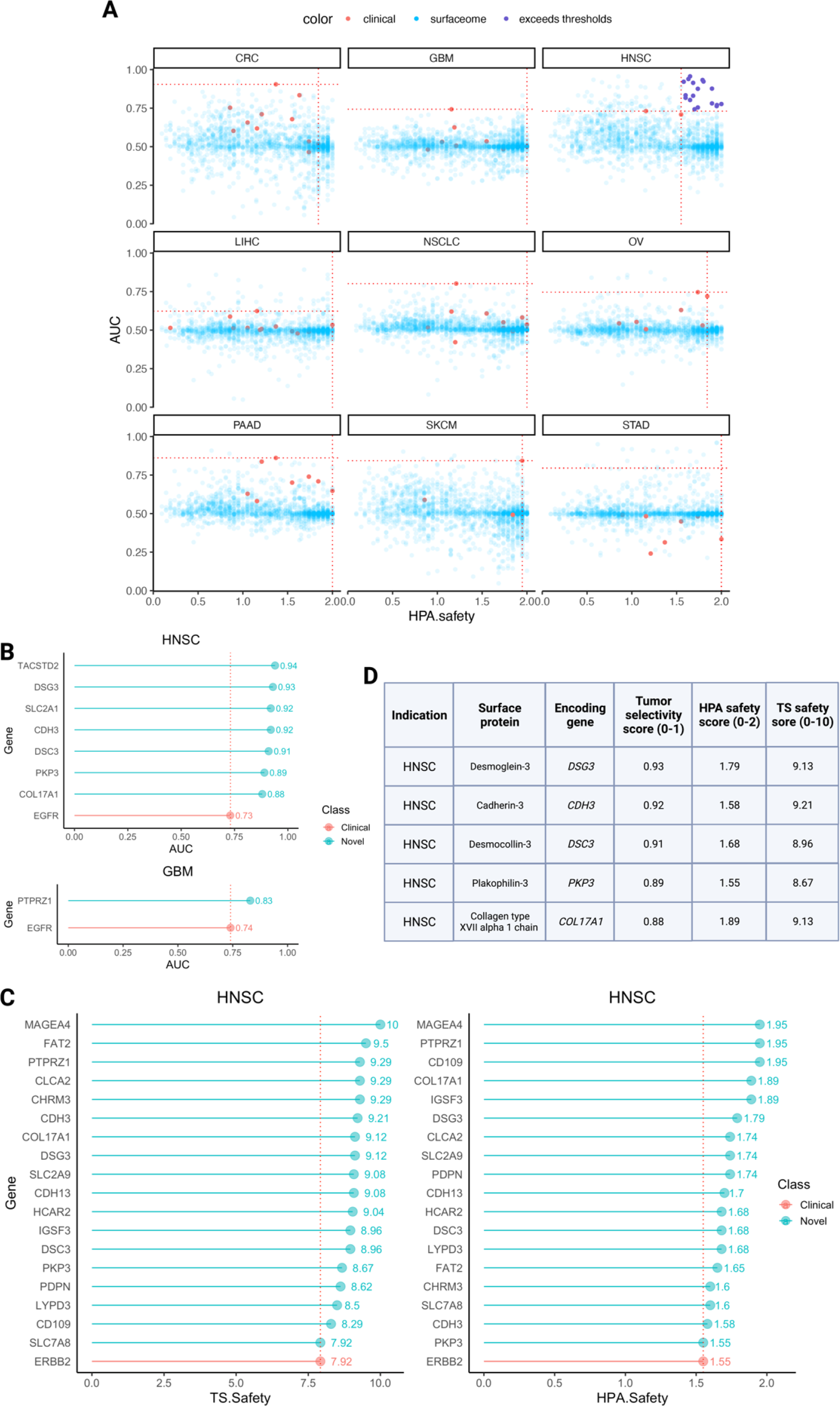
Identification of novel CAR targets with improved selectivity and safety compared to leading clinical CAR targets. **(A) 2D scatterplots of all surfaceome genes with HPA safety score on the x-axis and tumor selectivity score on the y-axis.** Red points represent clinical CAR target encoding genes for the respective cancer type. Horizontal thresholds denote the highest tumor selectivity score of clinical CAR targets pertaining to each cancer type, while vertical thresholds denote the highest respective HPA safety score of clinical CAR targets pertaining to the cancer type. Clinical CAR targets are shown in red, surfaceome genes which surpass both the horizontal and vertical thresholds (thus appearing in the upper right quadrant) are shown in dark purple. These points emerged only for HNSC. The remaining surfaceome genes are shown in transparent blue. Tabula sapiens safety scores were excluded for simplicity of visualization. **(B) Top selective (and yet safe) new targets.** Stems in red denote the leading clinical target by selectivity for each of the two cancer types, HNSC (left) and GBM (right), for which we found new selective targets. Stems in green denote the seven total surfaceome genes which emerged from our analysis as having greater tumor selectivity than the leading target in each cancer type. **(C) Top safe (and yet selective) new targets.** Stems colored red denote the leading clinical target by safety score, derived from HPA (left) and Tabula sapiens (right). Stems colored green denote the surfaceome genes which emerged from our analysis as having greater safety scores than the leading clinical CAR targets derived from each atlas. Both plots depict the same set of 18 newly suggested surfaceome genes for HNSC as well as the established target gene *ERBB2*. **(D) Top selective and safe new targets.** New surfaceome targets appearing as both highly selective and highly safe from the previous two iterations of the analysis (shown in **B & C**). Rows sorted by descending tumor selectivity scores.

We first aimed to identify novel *highly selective* CAR targets yet having potentially acceptable low toxicity. We searched for surface proteins that are encoded by genes whose selectivity scores across patients in a cancer type are significantly higher than the respective top currently established CAR target (employing a one-sided Wilcoxon signed rank test comparing patient-level selectivity scores, with FDR p-value < 0.1 denoting significance). To exclude targets with potentially extreme toxicity levels and only consider those that are most likely reasonable/safe, we applied lower bound thresholds to each of the sub-safety scores: 1.5 (out of 2) for HPA and 7.5 (out of 10) for Tabula sapiens. Notably, the only cancer types where we find new higher scoring selective targets are HPV-negative HNSC (seven targets) and glioblastoma (GBM; one target) (**Figure 4B & Additional File 2: Table S4**).

Second, studying a complementary perspective, we next searched for *highly safe targets*, that is those that have higher safety scores that the best approved/studied CAR target in a cancer type. We concomitantly required that these targets are at least moderately selective, with mean tumor selectivity scores ≥ 0.7. We required that new candidate targets safety scores surpass *both* atlases’ safety scores of the pertaining cancer-type-specific top target(s). Notably, here, we found highly targets only for HPV-negative HNSC. The eighteen newly suggested targets are shown in **Figure 4C** and **Additional File 2: Table S5** along with the established target *ERBB2*.

Finally, thirdly, we searched for target genes that have selectivity *and safety scores* that are both higher than those observed for the current targets studied *w*ithin each cancer type. We find five such targets, all for HPV-negative HNSC (**Figure 4D**). Indeed, charting the tumor selectivity scores and HPA safety scores of all surfaceome genes in two dimensions (Tabula sapiens safety scores were excluded to enable clearer visualization, but are provided in **Additional File 2: Table S3**), along with the cancer-type-specific thresholds for both metrics, revealed that HNSC is the only cancer type for which there exist any surface targets that surpass both thresholds and are thus more selective and safer than the leading existing CAR targets (**Figure 4A, 4D**).

### Further ranking CAR targets by their essentiality in tumor cells

To evaluate the potential of a surface protein to serve as a CAR target, it is of further interest to study its essentiality in cancer cells. A strong dependency on a target in its pertaining cancer type testifies to a biological dependency on that protein by the tumor cells and subsequently, points to a possible decreased potential for immune evasion via downregulation of the gene or loss of this antigen on the cell surface. To our knowledge, the dependency in cell lines of genes encoding current clinical CAR targets has not yet been studied; we thus analyzed the CRISPR DepMap [37] essentiality scores for these genes (**Figure 5A&B**, **Additional File 2: Table S6-7**) along with the untested candidate targets yielded by our pipeline (**Figure 5C**, **Additional File 2: Table S8**).

**Figure 5:**
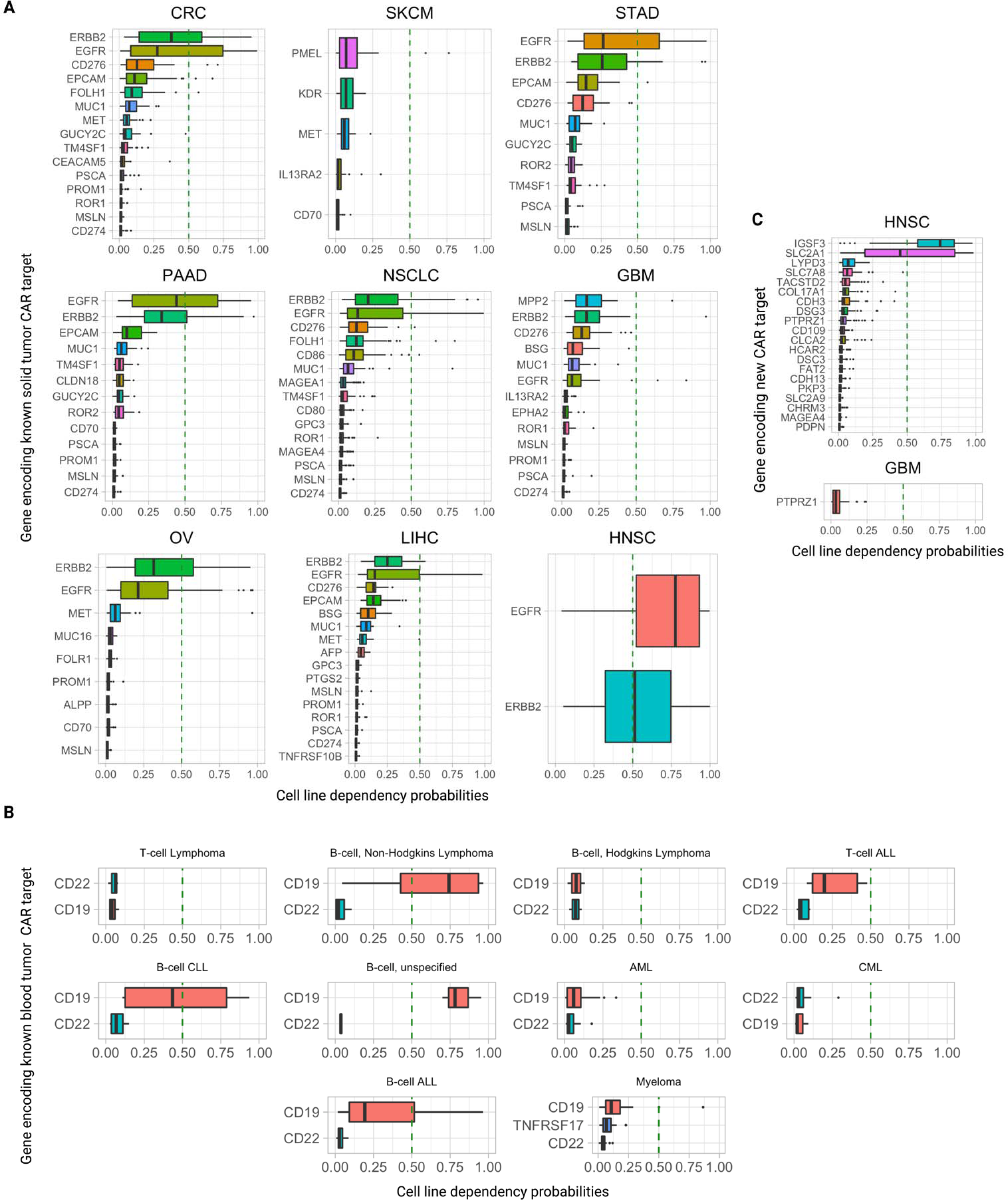
the dependency of genes encoding known CAR targets and our candidate new CAR targets in DepMap cell lines. **(A)** Probability of essentiality/dependency by cell lines on genes encoding existing solid tumor clinical CAR targets, for each of the nine cancer types studied. The vertical dashed line is at p(0.50), denoting the threshold above which a cell line is considered to be “dependent” on a gene. **(B)** Probability of essentiality/dependency by cell lines on genes encoding existing clinical liquid tumor CAR targets, for lymphomas, leukemias, and myeloma. Vertical dashed line is at p(0.50), denoting the threshold where a cell line is considered to be “dependent” on a gene. **(C)** Probability of essentiality/dependency by cell lines on genes encoding untested CAR targets we proposed for HNSC and GBM. Vertical dashed line is at p(0.50), denoting the threshold where a cell line is considered to be “dependent” on a gene.

To assess the essentiality of a given (cancer type, gene) pair, we analyzed the DepMap cell lines belonging to that indication (**Methods**) and considered a cell line dependent on that gene if *p*(dependence) ≥ 0.50, as defined by DepMap. For each target gene, we computed the mean *p*(dependence) over all cancer cell lines of that indication, and the fraction of cell lines in which the gene was deemed essential. We also note that DepMap subcategorizes ‘essential’ genes as either as ‘strongly selective’ or ‘common-essential.” The former denotes strong essentiality in a subset of cell lines and the latter denotes broad, pan-cancer essentiality in almost every cell line.

Intriguingly, most CAR targets in clinical trials exhibit quite low dependency scores across cell lines of their respective indications. One notable exception is the FDA-approved CAR-T target-encoded gene *CD19*, which is essential in 75% (9/12) of non-Hodgkin’s B-cell lymphoma cell lines, 25% (3/12) of B-cell acute lymphoblastic leukemia (B-ALL) cell lines, and 100% (3/3) of unspecified B-cell leukemia cell lines (**Figure 5B, Additional File 2: Table S6**). *CD19* is also deemed ‘strongly selective’ by DepMap. Interestingly, the other blood tumor clinical CAR-T target genes, *CD22* (in Phase 1 trials) and *TNFRSF17* (encoding the BCMA protein; FDA-approved against multiple myeloma), are not essential in *any* cell line of their respective indications analyzed here.

In solid tumors the emerging overall picture is similar. Almost all clinical CAR targets are not essential in any of the cell lines in the pertaining indications, with the striking exceptions of *EGFR* and *ERBB2,* which are highly essential across all their indications. For each cancer type they are indicated in, *EGFR* and *ERBB2* consistently rank the highest both in terms of fraction of cell lines they are essential in and the mean and median *p*(dependency) across cell lines (**Figure 5A**). *EGFR* is essential in 76% of HNSC cell lines and *ERBB2* is essential in 52% of HNSC cell lines (**Additional File 2: Table S7**). Out of the 35 solid tumor clinical CAR target genes that we collected and analyzed, *EGFR, ERBB2*, and *MET* are the only ones that are deemed essential by the CRISPR DepMap, all ‘strongly selective.’

Of the candidate CAR targets yielded by our pipeline, the vast majority are not essential in any cell lines of their respective cancer indication. However, two targets stand out: *IGSF3* and *SLC2A1*, which are essential in 80% and 45% of HNSC cell lines, respectively (**Figure 5C, Additional File 2: Table S8**). *IGSF3* is classified as ‘common-essential’ (*i.e.*, ‘pan-essential’) and *SLC2A1* is classified as ‘strongly selective’ by DepMap.

## Discussion

Overall, there are twenty unique genes encoding surface proteins that were identified by our pipeline across all three filtering configurations (top selective, top safe, and requiring both concomitantly). The twenty genes, the surface proteins they encode, and the families/classes of the respective proteins are shown in **Table 3**. Here we present a narrative summary of seven yielded by our analysis: the five superior targets that are both highly selective and safe, and two that are already existing targets of CAR therapies or antibody-drug conjugates in different indications. Analogous narratives for the remaining candidate surface targets can be found in **Additional File 1: Supp Text**.

**Table 3.**
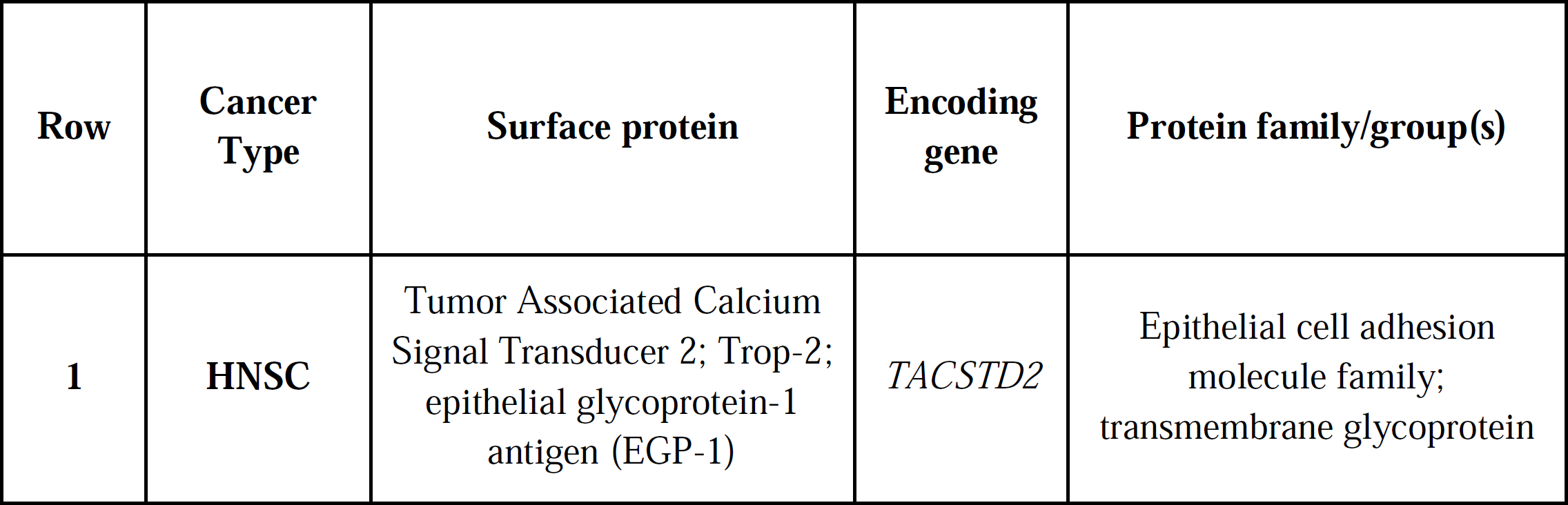
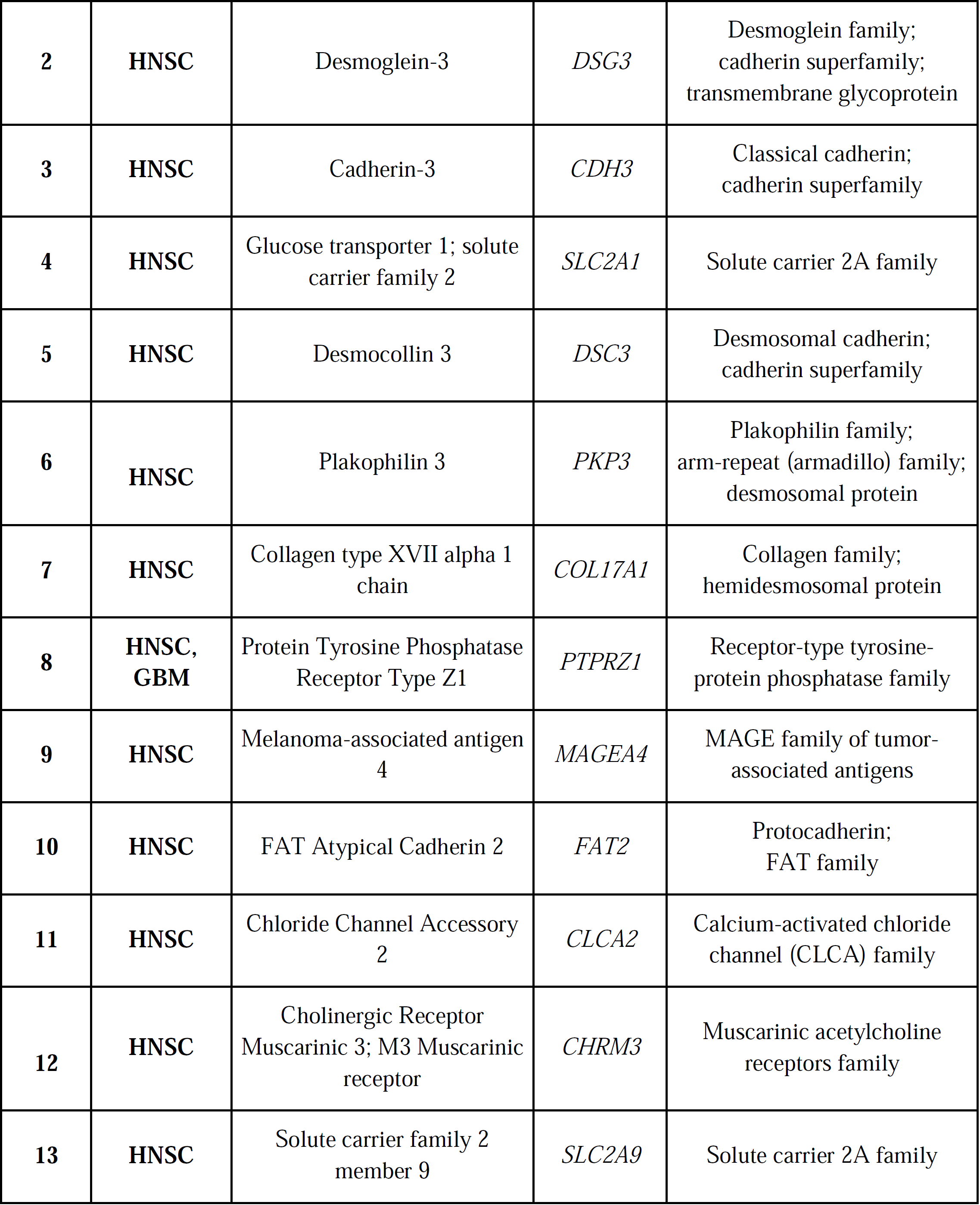
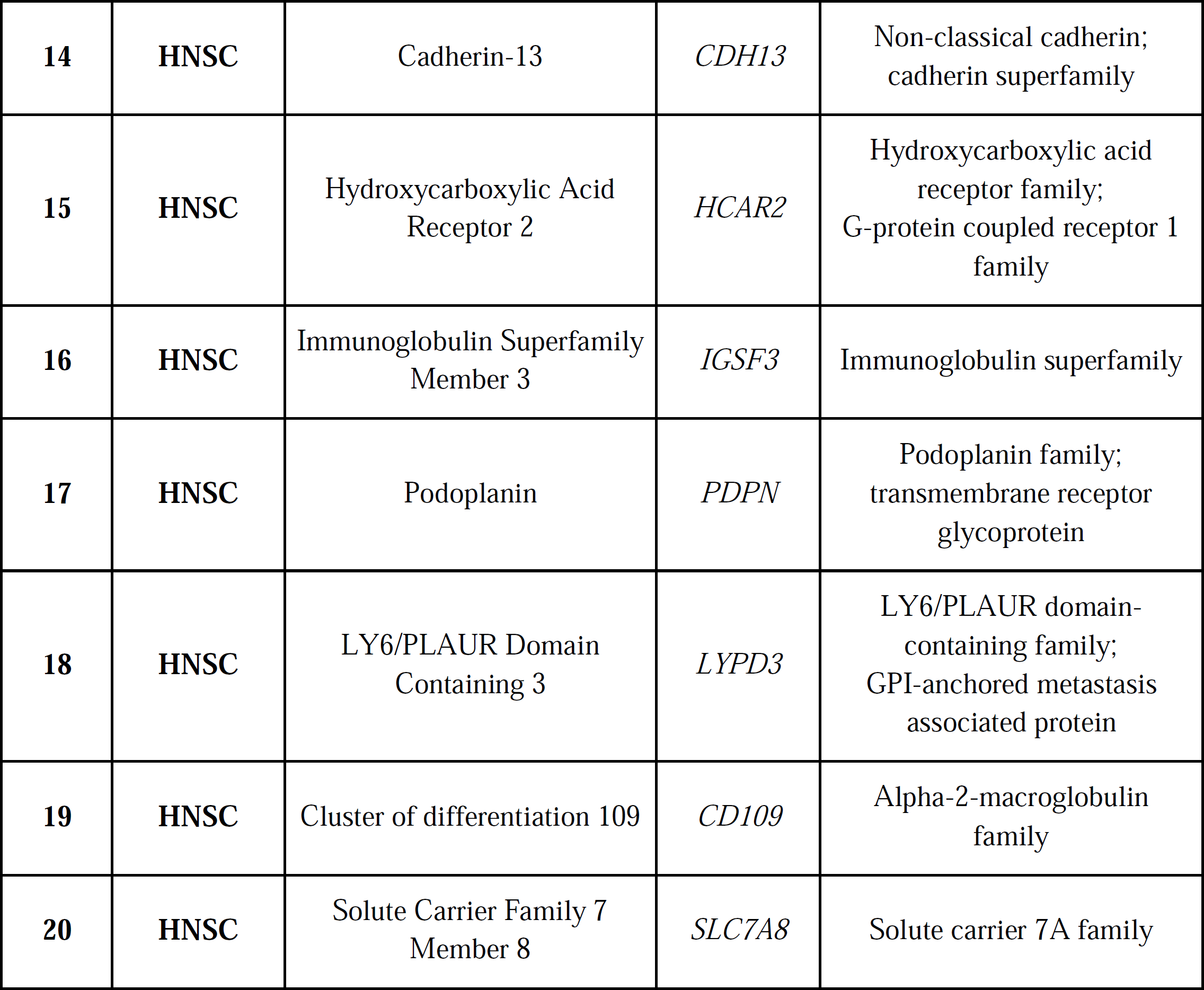
Families/classes of the twenty surface proteins encoded by genes resulting from our analysis as candidate CAR targets.

We begin with *TACSTD2*, having a very high selectivity score in HNSC (mean AUC=0.94, one-sided Wilcoxon signed rank test FDR p-value=0.065). It encodes the tumor associated calcium signal transducer 2, also known as “Trop-2” and “epithelial glycoprotein-1 antigen”, a calcium signal transducing cell surface receptor and known tumor-associated antigen. Among all cancers in The Cancer Genome Atlas (TCGA) [14], HNSC harbors the third highest median *TACSTD2* expression levels (265.2 FPKM). In the HPA Pathology Atlas [15], antibody staining revealed medium to high membranous and/or cytoplasmic positivity in two of four HNSC patients. Biallelic loss-of-function mutations in *TACSTD2* are known to cause gelatinous drop-like corneal dystrophy [16], which suggests that expression of this gene is functionally important only the eye. Notably, Trop-2 is the target of sacituzumab govitecan [17], an antibody-drug conjugate that is FDA-approved to treat metastatic triple-negative breast, HR-positive breast, and urothelial cancers. The existence of this approved drug may provide a basis for expedited anti-Trop2 CAR development for patients with HNSC and other solid tumors.

Next, *DSG3* both improved tumor selectivity (AUC=0.93, FDR p=0.065) *and* safety (Tabula sapiens score=9.13; HPA score=1.79) over the leading extant targets in HNSC. This gene encodes the transmembrane glycoprotein desmoglein-3, which belongs to both the desmoglein family and the cadherin cell adhesion molecule group. *DSG3* median gene expression levels were the highest in HNSC among all TCGA cancer types (median=155 FPKM), and its encoded protein was detected in two of four HNSC patients via membranous and/or cytoplasmic antibody staining in the HPA Pathology Atlas. Additionally, *DSG3* was previously found to be overexpressed at both the RNA and protein levels in head and neck cancer [18]. Inactivating mutations in the *DSG3* gene can lead to the formation of acantholytic blisters in the oral and laryngeal mucosa [19]. Importantly, the autoimmune disease pemphigus vulgaris is caused by autoantibodies targeting desmoglein-3 [20], resulting in blistering and erosions of the skin and mucosae (accordingly, desmoglein-3 is also known as “pemphigus vulgaris antigen”). Given this critical caveat, we caution that developing CARs against all DSG3-expressing cells is likely risky, and that a better strategy is likely to target only DSG3-overexpressing cells.

*CDH3* improved upon both the leading clinical tumor selectivity and safety scores in HNSC (AUC=0.92, FDR p-value=0.065; TB safety=9.21; HPA safety=1.58). It encodes the cadherin-3 protein, which is a member of the cadherin superfamily, cell adhesion proteins that are dependent on calcium and enable cells to stick together by binding to each other. *CDH3* exhibited the highest median RNA expression in HNSC among all cancer types in the TCGA (median=90.9 FPKM). Moreover, HPA Pathology antibody staining revealed all four out of four HNSC patients exhibited medium/high CDH3 protein expression. Previous studies have reported *CDH3* overexpression and association with poor prognosis in HNSC [21–23]. Biallelic mutations in *CDH3* can cause recessive congenital hypotrichosis with juvenile macular dystrophy [24] a rare disorder characterized by sparse hair and progressive vision loss, as well as ectodermal dysplasia, ectrodactyly, and macular dystrophy syndrome, similar to the former but additionally causing congenital limb malformation [25].

*DSC3*, another cadherin gene, improves upon both tumor selectivity and safety in HNSC (AUC=0.91, FDR p=0.065; TS safety=8.96; HPA safety=1.68) and encodes the desmocollin-3 protein. As a desmosomal protein, it is mostly present in epithelial cells and is essential for the formation and adherence of desmosome cell-cell junctions. In the HPA Pathology Atlas, *DSC3* is “cancer enhanced” in head and neck cancer and cervical cancer, with highest median expression in HNSC (94.5 FPKM) out of all cancer types. Moderate/strong cytoplasmic positivity was seen in HNSC, while DSC3 was generally lowly expressed in most other cancers [15]. While studies on this protein’s role in HNSC have been limited to the best of our knowledge, reduced expression of DSC3 in oral squamous carcinomas has been previously correlated with histological grade [26]. Biallelic inactivating mutations in *DSC3* cause skin blistering and hypotrichosis [27].

Next, *PKP3*, encoding the protein plakophilin-3, improves upon both the leading clinical tumor selectivity and safety scores for HNSC (AUC=0.89, FDR p=0.065; TB safety=8.67; HPA safety=1.55). It belongs to the plakophilin family of desmosomal proteins and links the desmosomal plaque to cytoskeletal intermediate filaments. While detected in many TCGA cancer types, it displayed highest median expression in HNSC (82.8 FPKM). Moderate to high cytoplasmic immunoreactivity of this protein was yielded by the majority of HNSC cancers stained in the HPA Pathology Atlas. Intriguingly, while the upregulation of *PKP3* has been observed during development of many cancer types, its potential tumor suppressive role was also reported in HNSC, with its expression prognostically favorable, denoting a paradoxical role of it and other desmosomal proteins in HNSC [28]. There are no known monogenic disorders caused by mutations in *PKP3*.

The next candidate target, *COL17A1*, encodes the collagen type XVII alpha 1 chain. This gene has high tumor selectivity and safety scores for HNSC (AUC=0.88, FDR p=0.084; TS safety=9.13; HPA safety=1.89). The encoded protein is part of hemidesmosomes, which are protein complexes mediating the adhesion of the dermis and epidermis. The HPA Pathology Atlas reports *COL17A1* as specifically “cancer enriched” in HNSC (164.9 FPKM), while its expression was detected in many cancer types. Moderate to strong cytoplasmic and membranous staining of the encoded protein was observed in HNSC samples. Collagen XVII has been found to frequently present in HNSCs, especially those with an oral cavity or larynx origin(29).

Biallelic inactivating mutations in *COL17A1* cause intermediate junctional epidermolysis bullosa 4 [30], a genetic disorder characterized by skin blisters and erosions, nail dystrophy, and alopecia. *COL17A1* mutations can also cause epithelial current erosion dystrophy, a genetic disorder characterized by corneal erosion [31].

*MAGEA4* has higher safety scores in HNSC (TS safety=10; HPA safety=1.95). It encodes the melanoma-associated antigen 4, which regulates cell proliferation by stopping cell cycle arrest at G1 phase and decreases apoptosis mediated by p53. Importantly, the MAGE-A4 protein is already an extant CAR-T target against non-small cell lung cancer (NSCLC) [10] but has not been studied in HNSC patients. Thus, repurposing CAR therapies targeting this protein to patients with MAGEA4+ HNSC may be low cost and high yield. HPA Pathology Atlas defines *MAGEA4* to be “cancer enhanced” in lung cancer in the TCGA, with comparable expression in head and neck cancer. Medium to high protein expression was observed in HNSC samples by antibody staining. Its widespread expression has been previously observed in a variety of head and neck cancers [32–36]. There are no known monogenic disorders caused by mutations in *MAGEA4*.

We additionally explored the *in vitro* dependency by tumors on genes encoding known CAR targets (to our knowledge, the first systematic interrogation) and our newly uncovered candidate CAR targets. The strong essentiality of *CD19* in a significant fraction of B-cell lymphomas and leukemias is unsurprising given that CD19 is a B-cell marker and has important function in B cell receptor signaling. However, other blood tumor clinical CAR-T target genes, *CD22* and *TNFRSF17*, are not essential in any cell line of their respective indications. In solid tumors, the high essentiality of *EGFR* and *ERBB2* may suggest that these genes are highly important to the viability of cancer cells in their respective solid tumor indications. The essentiality of *IGSF3* and *SLC2A1* in a significant fraction of HNSC cell lines may indicate their importance to the viability of HNSC tumor cells. Consequently, the ability of these cancer cells to downregulate *IGSF3* and *SLC2A1* to evade CAR cell killing might be limited.

In closing, we note that our findings are aligned with those reported in a recent study that has characterized the dependence of the genes constituting the surfaceome in cancer cell line growth and found that the percentage of surfaceome genes defined as essential in DepMap “[was] substantially less than” the percentage of non-surfaceome genes defined as essential; only 4.1% of the former were essential versus 14.0% of the latter [7]. We note, however, that DepMap quantified *in vitro* gene essentiality, whose relation to *in vivo* essentiality observed in patients’ tumors remains to be further elucidated.

The key finding of our study is a set of genes encoding cell surface proteins that may constitute promising new targets for engineering new CARs, mostly to treat HNSC and to a lesser extent GBM tumors. The targets we found have higher tumor selectivity and safety scores versus those of leading extant targets in clinics. Identified primarily by the analysis of tumors expression at the single cell level, most of our proposed targets have additional supportive evidence from human reference atlases, TCGA expression, pathology staining and previous gene-specific publications. Our analysis of dependency scores in the DepMap illuminates the intriguing phenomenon that the vast majority clinical CAR targets are not biologically essential to cancers, except for: *CD19, EGFR, ERBB2*, and *MET*. Of the targets yielded by our analysis, *IGSF3* and *SLC2A1* exhibited strong essentiality in HNSC cell lines.

Our study has a few limitations. First and foremost, we have used gene expression as a proxy for protein expression, but it is ultimately proteins that are targeted by CARs. When advances in high-throughput single-cell proteomics technology come to the forefront in the future, it will be pertinent to apply the approach presented in this study here to evaluate the discriminatory potential of membrane protein receptors as viable CAR targets. In the meantime, with the accumulating plethora of scRNA-seq datasets of tumors, gene expression is the best representation of cell genomics available to us. Second, the use of the AUC metric to evaluate target discrimination potential has limitations. In practice, optimal discrimination relies on knowing antigen detection thresholds of CAR-T cells. Currently, our control in setting antigen detection density of CARs is poor, but emerging efforts may allow us to tune antigen detection cutoffs^5^. This would allow us to evaluate CAR target viability more practically. Finally, in future studies, it would be of great interest to evaluate dual- and multi-targeted CARs, based on the approach outlined herewith. The computational framework enabling such combinatorial investigations has already been laid out in MadHitter [6], but for simplicity and practical applicability, we aimed to focus here on finding optimal single CAR surfaceome targets.

In summary, employing a systematic, data-driven analysis, we first chart the landscape of existing CAR targets in the clinic, identifying the leading targets in each cancer type based on tumor selectivity and safety metrics that we have computed. Next, from single cell transcriptomics data, we have performed a genome wide search across many different cancer types to identify new and candidate CAR targets with both better selective and safety scores than those of extant clinical targets. Remarkably, in almost all cancer types, we could not find such better targets, testifying to the near optimality of the current target space, at least in accordance with our measures. However, one striking exception is HPV-negative HNSC, for which there is a dearth of existing targets in clinical trials. Still, bearing that in mind, specifically, our investigation has discovered 20 new and untested CAR targets for treating HNSC and GBM tumors more precisely and safely than extant targets, which are each followed up with in a thorough literature survey and an analysis of their *in vitro* indication specific essentiality. Five of these 20 have selectivity and safety scores that are both higher than those computed for the existing CAR targets, whose further investigation may lead to better CAR treatments in HNSC.

## Additional files

### Additional File 1 (.txt)

**Supp Text:** Literature review on candidate targets (the fourteen not mentioned in the main text).

**Fig S1:** the distributions of safety scores in surfaceome genes versus indication-specific CAR targets. (A) shows HPA safety scores, (B) shows Tabula Sapiens safety scores.

**Fig S2:** the distributions of safety scores in indication-specific CAR targets versus top 10 tumor-selective surfaceome genes in each indication. Wilcoxon rank-sum test comparison p-values shown at bottom of each plot. (A) shows HPA safety scores, (B) shows Tabula Sapiens safety scores.

### Additional File 2 (.xlsx)

**Table S1:** (gene x cancer type) cells list all the patient-specific tumor selectivity scores (aka tumor-vs-nontumor classification AUCs) of each surfaceome gene in the respective cancer type. ‘N/A’ denotes genes that weren’t profiled for single cell datasets pertaining to the respective cancer type.

**Table S2:** (gene x tissue group) values denote levels of protein abundance measured in tissue group, such that: ND-L < Variable < M-H. Safety scores shown in rightmost column.

**Table S3:** All (gene x organ) cells contain fraction of cells expressing gene in respective organ. Overall safety score provided in rightmost column.

**Table S4:** Top selective (and yet safe) new targets. The first two rows (marked by *) denote the leading clinical targets by selectivity for the two indications, HNSC and GBM, for which we found new targets. The table is sorted by descending tumor selectivity scores.

**Table S5:** Top safe (and yet selective) new targets. The first row (marked by *) denotes the leading clinical target by safety for HNSC. The second row (marked by **) denotes a protein which is already a CAR target for the NSCLC indication (MacKay et al.). The table is sorted primarily by descending Tabula Sapiens safety scores, with ties broken in order of descending HPA safety scores.

**Table S6:** Dependency in cell lines of genes encoding CAR-T targets in blood tumors.

**Table S7:** Supplementary Table 7. Dependency in cell lines of genes encoding CAR-T targets in solid tumors.

**Table S8:** Supplementary Table 8. Dependency in cell lines of each candidate target gene yielded by our pipeline. Rows highlighted in green are genes that are classified as ‘strongly selective’ or ‘pan-essential’ by DepMap and are considerably more essential than all other genes in the Table.

### Ethics approval and consent to participate

Not applicable.

### Consent for publication

Not applicable.

## Availability of data and materials

All data analyzed in this study are publicly available via their respective cited sources. TISCH single-cell RNA-seq data were downloaded from http://tisch1.comp-genomics.org/. HPA data were downloaded from https://www.proteinatlas.org/about/download. Tabula sapiens data were downloaded from https://figshare.com/projects/Tabula_Sapiens/100973. CRISPR DepMap data were downloaded from https://figshare.com/articles/dataset/DepMap_22Q2_Public/19700056/2.

## Author’s contributions

AAS and ER supervised the study. SM, AAS, and ER conceived and designed the study. SM and SS analyzed data and designed the figures. SM, JSG, EEWC, AAS, and ER drafted and revised the manuscript. All authors read and approved the final manuscript.

## Acknowledgements

The NIH HPC Biowulf cluster (http://hpc.nih.gov) was used for computational analyses.

