## Additional File 1 (Supplementary Text and Figures) for "Pan-cancer analysis of patient tumor single-cell transcriptomes identifies promising selective and safe CAR targets in head and neck cancer"

**Additional File 1: Supplementary Text & Figures**

**Supp Text: Literature review on candidate targets (the fourteen not mentioned in the main text)**

In the main text, we presented 20 CAR targets for HNSC and described the seven most selective in some detail. Here, we describe the other 13 (= 20 -7) target genes and one target gene for GBM. References cited by number refer to the reference list at the end of this document not to the reference list in the main document.

The next candidate, *SLC2A1*, encodes the glucose transporter 1 protein, also known as solute carrier family 2 member 1 or GLUT1. It and improved upon the leading clinical HNSC tumor selectivity score (AUC=0.92, FDR p-value=0.065). The encoded cell surface protein plays a key role in transporting glucose through the blood-brain barrier in mammals. While its RNA expression was detected in all TCGA cancer types, it exhibited the highest median expression in HNSC (FPKM=202). In the HPA Pathology Atlas, moderate to strong cytoplasmic positivity of the protein was seen in most cancers, including HNSC. A previous report described *SLC2A1* as “one of the pivotal genes in cancer glycometabolism,” and found its elevated expression in TCGA tumor vs. GTEx normal controls in 22 cancer types, HNSC included [1]. The role of this gene has also been reported in oral squamous cell carcinoma [2]. Heterozygous mutations in this gene can cause GLUT1 deficiency syndrome 1, a metabolic disorder characterized by seizures, developmental delay, and microcephaly [3], and the similar but milder GLUT1 deficiency syndrome 2 [4]. In DrugBank [5], *SLC2A1* is listed as a target of a variety of approved, investigational, and experimental compounds, including etomidate, glucosamine, and ascorbic acid [6–8].

*FAT2* has higher safety scores in HNSC, encoding FAT atypical cadherin 2 (TS safety=9.5; HPA safety=1.65). The HPA Pathology Atlas defines the *FAT2* gene to be “group enriched” in cervical cancer, head and neck cancer, lung cancer, and urothelial cancer, with highest median TCGA RNA expression in HNSC (18.8 FPKM) among all cancer types. Antibody staining yielded cytoplasmic and membranous staining in only urothelial cancers, while all other cancer types were negative. Not much is currently known about the role of *FAT2* in cancer, however, targeting the paralog *FAT1* was previously shown to be beneficial in treating oral squamous cell carcinoma, a type of HNSC [9] Mutations in *FAT2* are associated with autosomal dominant spinocerebellar ataxia type 45 [10].

*CLCA2*, encoding the chloride channel accessory 2 protein, has higher safety scores of HNSC (TS safety= 9.29, HPA=1.74). This protein modulates the flow of chloride ions across the cell membrane in a calcium-dependent manner and is involved in cell adhesion, particularly that of basal cells, and layering of squamous cells. The HPA Pathology Atlas reports *CLCA2* to be “cancer enhanced” in head and neck cancer in the TCGA, along with strong cytoplasmic and membranous staining in HNSC samples. Previously, this gene was reported to be overexpressed in HNSC and its expression levels were predictive of sensitivity to EGFR inhibitors in cell lines [11]. Mutations in *CLCA2* are not associated with any known monogenic diseases.

*CHRM3* also has higher safety scores in HNSC (TS safety=9.29; HPA safety=1.6), encoding cholinergic receptor muscarinic 3, also known as M3 muscarinic receptor. This receptor is a G protein-coupled receptor and triggers a variety of cellular responses, such as reduction of adenylate cyclase activity, breaking down of phosphoinositides, and control of potassium channels through G protein action. The HPA Pathology Atlas finds low cancer type specificity of *CHRM3* gene expression in TCGA, but its median expression ranks second highest in HNSC vs. all other cancer types (median=1 FPKM). *CHRM3* has been observed to be highly expressed in HNSC cells [12]. Seventy-five drugs are listed in DrugBank for which *CHRM3* is a reported target. Many drugs target muscarinic receptors, including anticholinergic drugs (*e.g.*, tiotropium, ipratropium) and cholinergic agonists (*e.g.*, pilocarpine). These drugs do not specifically target CHRM3, but affect all muscarinic receptors, including M3 [5]. In one family, a homozygous mutation in *CHRM3* was associated with prune belly syndrome, a type of urinary bladder disease [13].

*SLC2A9* has improved safety scores in HNSC (TS safety=9.08; HPA safety=1.74). This gene encodes the solute carrier family 2 member 9 protein, also known as GLUT9, and is a part of the family of SLC2A facilitative glucose transporters. Its former name GLUT9 is misleading because the primary transport function of GLUT9 is to transport uric acid and it may also transport fructose [14]. Biallelic germline mutations of GLUT9 cause the rare disorder hypouricemia [15]. This disorder may be considered as a conceptual opposite of the much more common disorder gout that is characterized by excess uric acid. Consequently, the drug lesinurad, which inhibits GLUT9 and other transporters [16], has been FDA-approved to treat gout. Another approved gout treatment, allopurinol, was found (long after its approval) to inhibit GLUT9 [17]. The wide usage of these gout drugs suggests that GLUT9 inhibition is safe in many individuals although there can be serious side-effects [18] and adverse germline-drug indications are known [19]. Furthermore, a chemical found naturally in tea has been found to inhibit GLUT9 in rats [20]. Additionally, in the DrugBank, GLUT9 is listed as a target for a variety of approved and investigational compounds, including probenecid, losartan, uric acid, fludeoxyglucose (18F), dextrose, D-glucose, and olsalazine [5]. Because of the side effects of GLUT9 inhibitor drugs, we caution that an unmodulated CAR strategy of killing all GLUT9-expressing cells is likely unsafe, but if one could target only GLUT9-overexpressing cells, this gene is a promising HNSC target. Accordingly, we observe that *SLC2A9* RNA expression is detected in many TCGA cancer types, HNSC ranked second highest for its median expression (2.8 FPKM). HPA Pathology cytoplasmic staining of the protein was positive in HNSC, among most cancers.

Next improving on safety scores for HNSC was *CDH13*, encoding the Cadherin 13 protein (TS safety=9.08; HPA safety=1.70). CDH13 is a nonclassical member of the cadherin family. It plays a role in the differentiation and growth of neurons and exhibits hypermethylation across a variety of cancer types (<https://www.ncbi.nlm.nih.gov/gene/1012>). Its median TCGA RNA expression levels were highest in HNSC (4.4 FPKM) out of all cancer types. However, most cancers in the HPA Pathology Atlas, including HNSC, displayed negative antibody staining of the protein. *CDH13* hypermethylation has been previously observed in HNSC cell lines [21]. Mutations in *CDH13* are not associated with any known monogenic diseases.

*HCAR2* has improved safety scores in HNSC (TS safety=9.04; HPA safety=1.68). The gene encodes the hydroxycarboxylic acid receptor 2 (also known as “niacin receptor 1” and GPR109A) and is a G-protein coupled receptor. When activated, this protein suppresses fat breakdown and atherosclerosis formation and causes blood vessel dilation. The HPA defines this gene as “group enriched” in cervical cancer, head and neck cancer, lung cancer and urothelial cancer, with its highest median RNA expression in HNSC (13.2 FPKM) out of all cancer types. Cytoplasmic and membranous immunoreactivity was seen in an HNSC patient [22]. To the best of our knowledge, not much has been previously reported in the literature about the role of *HCAR2* in HNSC. HCAR2 is targeted by niacin (vitamin B3), which is used to treat high cholesterol and niacin deficiency [5]. Mutations in *HCAR2* are not associated with any known monogenic diseases.

*IGSF3*, encoding the immunoglobulin superfamily member 3, also has improved safety scores in HNSC (TS safety=8.96; HPA safety=1.89). This protein’s structure is similar to that of immunoglobulin, and it possesses multiple domains resembling V-type Ig-like domains. Although the HPA Pathology Atlas finds low cancer specificity of its RNA expression in the TCGA, its median expression is second highest in head and neck cancer (25.1 FPKM), following melanoma (37.6 FPKM). Protein expression via antibody staining was observed in a HNSC patient [22]. Although the specific role of this gene in HNSC has not previously been explored to our knowledge, a prior study analyzing the immunoglobulin superfamily interactome uncovered several hundred new receptor-ligand interactions, cell-type-specific interactions, and dysregulation of receptor-ligand crosstalk in cancers [23]. A homozygous deletion in the *IGSF3* gene was reported in a single family with bilateral lacrimal duct obstruction [24].

*PDPN,* which encodes podoplanin (Tabula safety= 8.63; HPA safety=1.74) has also improved safety scores. It plays a role in a variety of cellular processes, including cell migration and adhesion, formation of lymphatic vessels, platelet aggregation, tumor progression, and proliferation of normal lung cells. The HPA Pathology Atlas defines *PDPN* as “group enriched” in glioma, head and neck cancer, and testis cancer in the TCGA RNA expression. Cytoplasmic and membranous staining of podoplanin was observed in an HNSC patient [22]. Podoplanin has been previously shown to play a role in HNSC prognosis [25] and its knockdown inhibited nasopharyngeal carcinoma proliferation, migration, and invasion [26]. Mutations in *PDPN* are not associated with any known monogenic diseases.

*LYPD3* is another target with improved safety scores of HNSC (TS safety=8.5; HPA safety=1.68). This gene encodes LY6/PLAUR domain containing 3, found in extracellular space. It assists in cell migration and is believed to enable binding with laminin and play a role in cell-matrix adhesion. *LYPD3* is denoted by the HPA as “cancer enhanced” in cervical cancer and head and neck cancer in TCGA RNA expression, with its highest median expression in HNSC (198.1 FPKM). Strong cytoplasmic and membranous staining of the protein was detected in all four HNSC patients in the HPA Pathology Atlas. LYPD3 has been previously studied as a target for antibody-drug conjugates (ADCs) in treating squamous cell carcinomas of varying origins, including HNSC [27]. *LYPD3* expression has also been inversely associated with clinical outcome (patient death) in head and neck cancer [28]. Mutations in *LYPD3* are not associated with any known monogenic diseases.

An additional improved HNSC safety scoring target is *CD109* (TS safety=8.29; HPA safety=1.95), encoding the cluster of differentiation 109 molecule. CD109 is a glycoprotein anchored to the cell surface via a GPI. CD109 is found on the surface of platelets, activated T-cells, and endothelial cells, and binds to and inhibits TGF-beta signaling. Although its RNA expression was detected in all TCGA cancer types, HNSC ranked highest by median expression level (13.4 FPKM). Protein expression was not detected in most cancer tissues via antibody staining, including HNSC [22]. CD109 expression has been linked to unfavorable prognosis in oropharyngeal squamous cell carcinomas [29], and high expression of CD109 has been reported in various squamous cell carcinomas, including HNSC [30]. Mutations in *CD109* are not associated with any known monogenic diseases.

The final HNSC target emerging from our analysis with improved safety scores in HNSC targets (TS safety=7.92; HPA safety=1.6) is *SLC7A8*, encoding the protein solute carrier family 7 member 8, also known as L-type amino acid transporter 2 (LAT2). LAT2 enables transport of neutral amino acids and thyroid hormones across the plasma membrane. While detected in all TCGA cancer types by RNA expression, its median expression levels were highest in HNSC (26.9 FPKM). In the HPA Pathology Atlas, weak cytoplasmic/membranous staining was detected in three out of four HNSC patients. Increased LAT2 expression has been reported previously in HNSC tumors [31]. Mutations in *LAT2* are not associated with any known monogenic diseases.

Beyond HNSC, *PTPRZ1* has higher selectivity scores than extant targets in GBM (AUC=0.83, FDR p=5.53x10^-5^) and higher safety score in HNSC (TB safety=9.29; HPA safety=1.95). *PTPRZ1* encodes the protein tyrosine phosphatase receptor type Z1, which negatively regulates the proliferation of oligodendrocyte precursor cells in the spinal cord and is essential for their differentiation into mature oligodendrocytes. It may also stop oligodendrocytes from undergoing apoptosis and dephosphorylate signaling proteins to establish contextual memory. The HPA Pathology Atlas found *PTPRZ1* to be “cancer enriched” in glioma (median=157.6 FPKM), markedly higher than its expression in all other cancer types in the TCGA. Median expression was second highest in HNSC (9.3 FPKM). Moreover, gliomas exhibited moderate to strong nuclear and cytoplasmic positivity of the protein via antibody staining. Previous studies reported an association between *PTPRZ1* expression and stem cell-like properties [32], and its inhibition by a small molecule abrogated stemness and tumorigenicity of GBM cells [33]. *PTPRZ1* gene and protein expression has previously been shown to be abnormal and prognostic in oral squamous carcinomas [34,35]. Mutations in *PTPRZ1* are not associated with any known monogenic diseases.

**Supplementary Figures**

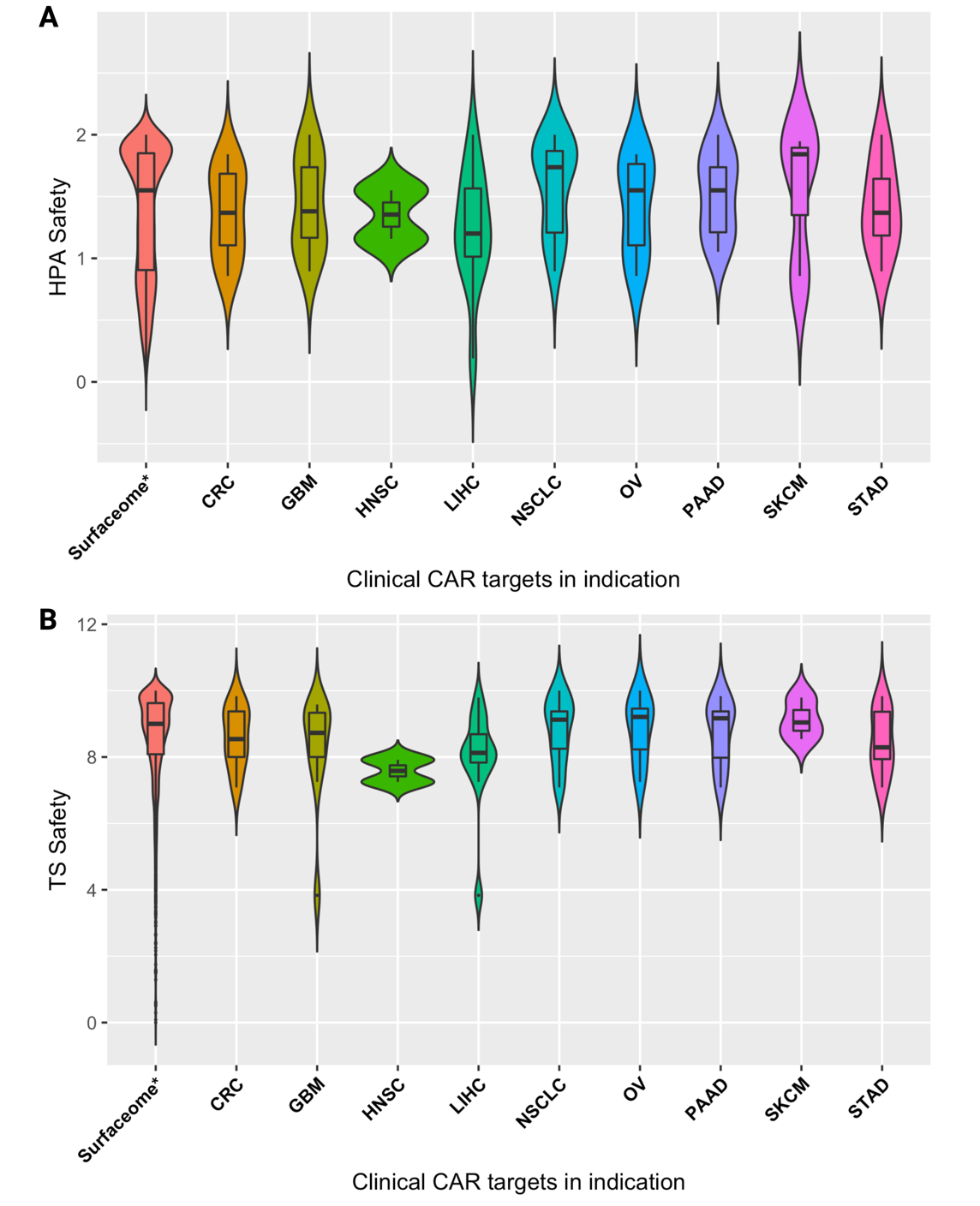

**Figure S1: the distributions of safety scores in surfaceome genes versus indication-specific CAR targets. (A)** shows HPA safety scores, **(B)** shows Tabula Sapiens safety scores.

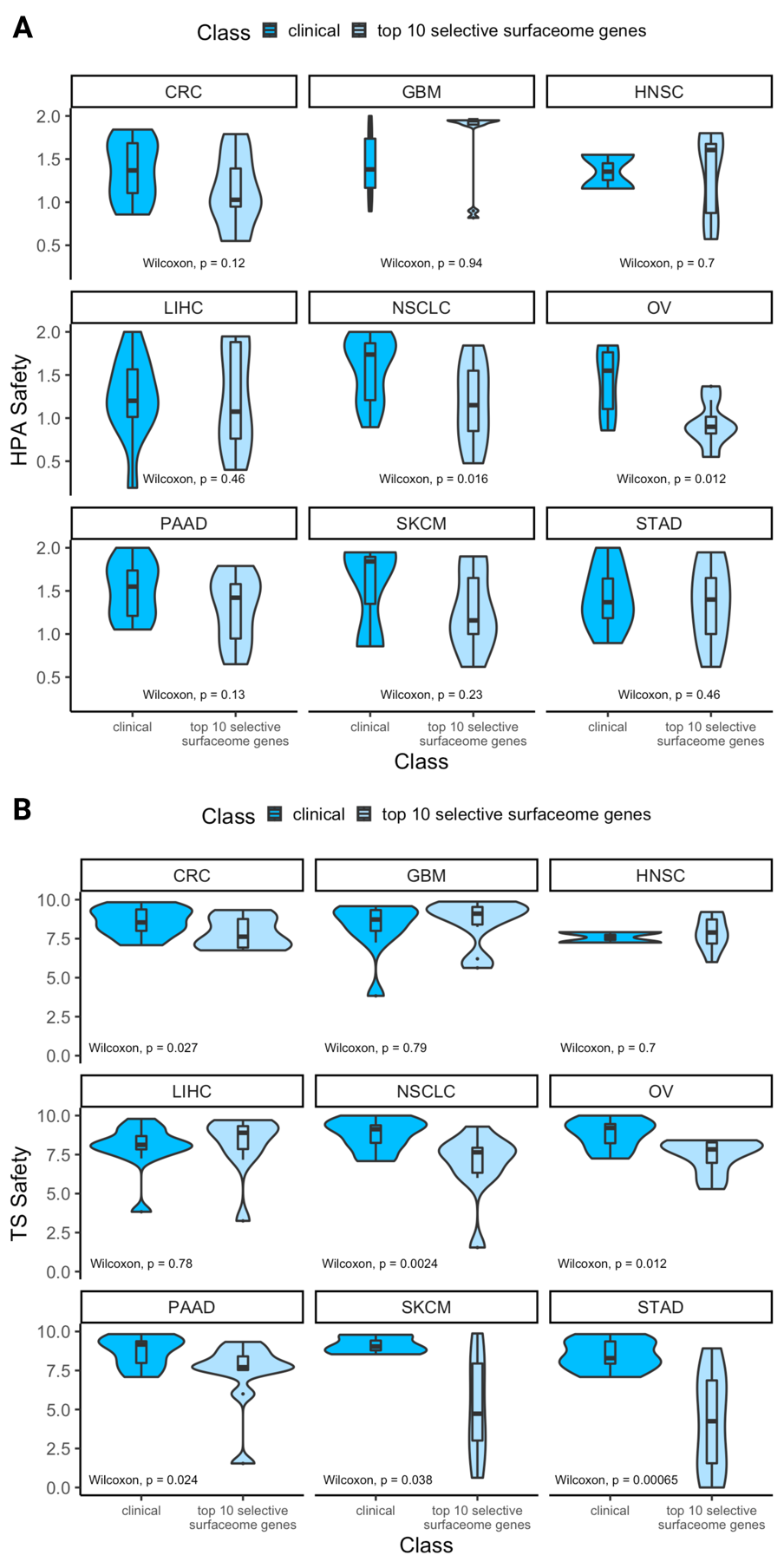

**Figure S2: the distributions of safety scores in indication-specific CAR targets versus top 10 tumor-selective surfaceome genes in each indication.** Wilcoxon rank-sum test comparison p-values shown at bottom of each plot. **(A)** shows HPA safety scores, **(B)** shows Tabula Sapiens safety scores.
